# Influenza A virus NS1 protein binds as a dimer to the RNA-free PABP1 but not to the PABP1•Poly(A) RNA Complex

**DOI:** 10.1101/2020.08.10.245225

**Authors:** Cyrus M de Rozières, Simpson Joseph

**Affiliations:** Department of Chemistry and Biochemistry, University of California, San Diego, La Jolla, CA 92093-0314, USA

## Abstract

Influenza A virus (IAV) is a highly contagious human pathogen responsible for nearly half a million deaths each year. Non-structural protein 1 (NS1) is a crucial protein expressed by IAV to evade the host immune system. Additionally, NS1 has been proposed to stimulate translation because of its ability to bind poly(A) binding protein 1 (PABP1) and eukaryotic initiation factor 4G (eIF4G). We analyzed the interaction of NS1 with PABP1 using quantitative techniques. Our studies show that NS1 binds as a homodimer to PABP1, and this interaction is conserved across different IAV strains. Unexpectedly, NS1 does not bind to PABP1 that is bound to poly(A) RNA. Instead, NS1 only binds to PABP1 free of RNA, suggesting that translation stimulation does not occur by NS1 interacting with the PABP1 molecule attached to the mRNA 3’-poly(A) tail. We propose that NS1 binds to the eIF4G complex at the 5’-end of the mRNA and recruits the RNA-free PABP1, which may stimulate translation initiation by promoting the association of the ribosomal subunits.

## Introduction

Seasonal Influenza A Virus (IAV) infection causes tens of thousands of deaths each year and billions of dollars lost in productivity with potential for greater severity during epidemics and pandemics.^1^ IAV is a negative sense, single-stranded RNA virus that infects the lung’s epithelial cells causing acute respiratory distress upon infection.^2^ The IAV genome is made up of 8 different segments that code for roughly a dozen proteins required for successful infection and replication.^3–6^ One of these viral proteins, Non-structural protein 1 (NS1), is responsible for a multitude of functions to help IAV proliferation, including the downregulation of the innate immune response, regulation of specific signaling pathways, and selective translation.^7^ NS1 is a 26 kDa protein made up of an RNA-Binding domain (RBD), a linker region, an effector domain (ED), and a C-terminal tail.^8^ NS1 is one of the most highly expressed proteins during IAV infection and yet does not get packaged into new viral particles, emphasizing its importance as a cell regulator during IAV infection. Studies of this protein and its roles have unveiled over a dozen different interactions with both host and viral proteins essential for successful viral replication.^7^

One of these interactions known to exist but not yet well understood is with the eukaryotic cytoplasmic Poly (A) Binding protein 1 (PABP1).^9^ PABP1 is a 72 kDa protein made up of four RNA recognition motifs, a homodimerization domain, and the PABC domain at the C terminal end.^10^ PABP1 is one of the most abundant proteins in the cell (∼4 µM) and is primarily responsible for stimulating translation initiation by binding to the poly(A) tail at the 3’ end of the mRNA.^11^ PABP1 bound to the 3’ poly(A) tail both protects the mRNA from exonucleases as well as stimulates translation initiation by interacting with eukaryotic initiation factors at the 5’ end of the mRNA.^12^ While it is unsurprising for viruses to target translation factors to replicate successfully, little is yet known about how IAV makes use of PABP1 during infection.

To date, what is known about the NS1•PABP1 interaction is that NS1 binds to PABP1 with a high affinity (K_D_ = 20 nM) and primarily involves the RBD of NS1 and the homodimerization domain of PABP1.^9,13^ Furthermore, NS1 cannot bind to both RNA and PABP1 simultaneously, suggesting a function that is independent of NS1’s other roles as an RNA binder.^13^ An interesting facet of this interaction is that the PABP1 homodimerization domain is long (∼170 residues) and predicted to be intrinsically disordered. This domain is primarily responsible for PABP1’s ability to multimerize on a poly(A) tail.^14^ Neither the purpose of this interaction nor the nature of the binding is well understood. We believe that further characterization of the interaction between NS1 and PABP1 is critical to understanding IAV infection. Additionally, identifying the binding mode can help design inhibitors that can specifically target this interaction to inhibit IAV infections. Here, we employed fluorescence polarization and gel-shift assays to study the binding of NS1 to PABP1. Our studies show that NS1 binds as a homodimer to PABP1, and the interaction is primarily electrostatic. Interestingly, NS1 binds to RNA-free PABP1 but does not bind to the poly(A)•PABP1 complex. Since NS1 also binds to eIF4G, a factor that is essential for translation initiation, the NS1•PABP1•eIF4G complex at the 5’-end of the mRNA may play a role in enhancing translation by promoting the recruitment of the ribosomal subunits.^15–18^

## Materials and Methods

### Purification of Recombinant NS1

H1N1 WSN wtNS1, H5N1 Nigeria wtNS1, H3N2 Udorn wtNS1, H3N2 Udorn WT NS1-RBD and NS1-RBD mutants were LIC cloned into the pETHSUL vector.^19^ NS1-RBD mutants were created by site-directed mutagenesis by either insertion of a C-terminal cysteine to make NS1-RBD FL or changing R35 and R46 to alanine to make MUT NS1-RBD. pETHSUL-NS1 constructs were transformed into *E. coli* BL21(DE3) cells. 2 L of cells were grown at 37 °C in LB/ampicillin to an OD_600_ of 0.6−0.8, and then induced with 1 M isopropyl β-D-1-thiogalactopyranoside for 2.5 h. Cells were pelleted, flash-frozen, and stored at −80 °C.

For H1N1 wtNS1 and H3N2 wtNS1 cells were resuspended in lysis buffer [25 mM Tris (pH 7.5), 500 mM NaCl, 5% glycerol, 0.5 mM EDTA, 1 mM PMSF, 0.1% Triton X-100, 8 mM dithiothreitol and 5 mM imidazole] and disrupted by sonication. The cell lysate was centrifuged at 20000g for 45 min at 4 °C. The supernatant was incubated with 4 mL of Ni-NTA beads for 15 min at 4 °C on a rotator. The slurry was poured over a column and washed with 50 mL of wash buffer [25 mM Tris (pH 7.5), 500 mM NaCl, 5% glycerol, 0.5 mM EDTA, 1 M KCl, 8mM dithiothreitol (DTT) and 50 mM imidazole]. Protein was eluted with elution buffer [25 mM Tris (pH 7.5), 500 mM NaCl, 5% glycerol, 8 mM DTT, 0.5 mM EDTA, and 250 mM imidazole]. Elutions were collected and concentrated using a 10K molecular weight cutoff (MWCO) concentrator until the volume was 5 mL. Protein was dialyzed at 4 °C overnight with constant stirring in storage buffer [25 mM Tris (pH 8.0), 100 mM NaCl, 10% glycerol, and 1mM TCEP-HCl] with Ulp1 protease (1:100) to remove SUMO tag. After dialysis the protein was incubated with 2 mL of Ni-NTA beads for 15 min at 4 °C on a rotator. The slurry was poured over a column and washed with storage buffer. Fractions were collected and analyzed by 12% sodium dodecyl sulfate−polyacrylamide gel electrophoresis (SDS−PAGE). Fractions were pooled and concentrated using a 3K MWCO concentrator, aliquoted, and flash-frozen. Concentrations of purified proteins were determined by the Bradford assay (Bio-Rad). Note that yields for these proteins post cleavage are low.

For H5N1 wtNS1 and H3N2 WT NS1-RBD and RBD mutants, cells were resuspended in lysis buffer [25 mM Tris (pH 7.5), 500 mM NaCl, 5% glycerol, 1 mM PMSF, 0.1% Triton X-100, 1 mM DTT and 20 mM imidazole] and disrupted by sonication. The cell lysate was centrifuged at 20000g for 45 min at 4 °C. Cell lysate was injected into FPLC and run over two HisTrap FF Crude 1 mL columns (Sigma-Aldrich) with FPLC running buffer [25 mM Tris (pH 7.5), 5% glycerol, and 0.5 mM TCEP-HCl]. Columns were washed with FPLC running buffer + 1 M NaCl and proteins were eluted with a step gradient (3%, 50%, 100%) of FPLC running buffer + 1 M Imidazole. Elutions were collected and run over a HiTrap Q 5 mL column (Sigma-Aldrich) with FPLC running buffer. A linear gradient from 0% to 100% FPLC running buffer + 1 M NaCl was used to elute proteins off of the column. Elutions were collected and analyzed by 16% Tricine−PAGE and A_280_/A_260_ measurements. Elutions free of nucleic acids were collected and concentrated using a 10K MWCO concentrator until the volume was 5 mL. Protein was incubated at 4 °C overnight on a rotator with Ulp1 protease (1:100) to remove SUMO tag. Protein post cleavage was injected into FPLC and run over two HisTrap FF Crude 1 mL columns (Sigma-Aldrich) with FPLC running buffer. The flow through fractions were pooled, concentrated and buffer exchanged with NS1-RBD Buffer [25 mM Tris (pH 7.5), 150 mM NaCl, 5% glycerol, and 0.5 mM TCEP-HCl] using a 3K MWCO concentrator, aliquoted, and flash-frozen. Concentrations of purified proteins were determined by the Bradford assay (Bio-Rad).

### Tryptophan Polarization Assay

Polarization studies were performed by using 2 μM of WT NS1-RBD or MUT NS1-RBD in NS1-RBD Buffer. Each sample (200 μL final volume) was transferred to a quartz cuvette and placed in a fluorometer (Jasco FP-8500) with a xenon lamp. The samples were excited at 295 nm, and the fluorescence emission intensity was measured at 350 nm. The excitation and emission bandwidth were set to 5 nm. A measurement of just NS1-RBD Buffer served as background. Three independent experiments were performed with three batches of protein.

### RNAs for Fluorescence Anisotropy

The poly(A)_18_ and ssCR1 (5′-GCUAUCCAGAUUCUGAUU-3′) RNA with a fluorescein dye attached to the 3′-end was purchased from GE Dharmacon. The dsRK1 with a fluorescein dye attached to the 5′-end was synthesized as two complementary RNAs: 5′-FL-CCAUCCUCUACAGGCG-3′ and 5′-FL-CGCCUGUAGAGGAUGG-3′. The S17 with a fluorescein dye attached to the 3′-end was synthesized as 5’-GGGTGACAGTCCTGTTT-FL-3’. All RNAs were deprotected and purified by denaturing urea−PAGE. All RNAs were resuspended in water, and their concentrations were determined by measuring the absorbance at 260 nm. All RNAs were stored at −80 °C in small aliquots. To make double-stranded RNA, equimolar amounts of sense and antisense RNAs were heated to 95 °C in 50 mM Tris (pH 8), 50 mM KCl, and 1 mM DTT for 2 min and then allowed to cool slowly to room temperature. The dsRNA was then aliquoted and stored frozen at −80 °C.

### Electrophoretic Mobility Shift Assay (EMSA)

An EMSA was performed by incubating proteins with Cy5 or fluorescein-labeled molecules [final concentrations of 100 nM] in anisotropy buffer to a final volume of 11 μL at room temperature for 1 h. After incubation, 1.3 μL of ice-cold 50% glycerol was added to the mix. The complexes were separated from unbound species by electrophoresis on a 0.7% nondenaturing gel using a SEAKEM GTG agarose solution (Lonza) made with 1× TBE buffer. Samples were separated at 4 °C in 1× TBE buffer for 1.5 h at a 66 V constant voltage. The gels were visualized by scanning with a FLA9500 Typhoon instrument using the Cy2 or Cy5 excitation laser at a 600 PMT voltage and 50 μm resolution. In cases where concentrations are not explicit, 500 nM PABP to 100 nM poly(A) RNA ratio was used with 5 μM NS1 when excess NS1 was used. Gels were analyzed and quantified with ImageJ software.^20^

### Fluorescence Anisotropy

Fluorescence anisotropy studies were performed using a fixed concentration of fluorescein-labeled RNA and an increasing concentration of protein in anisotropy buffer [50 mM Tris (pH 8), 50 mM KCl, 50 ng/μL *E. coli* total tRNA, 1 mM DTT, and 0.01% Tween 20].^21^ Note that the 50 mM KCl in the anisotropy buffer was exchanged for varying concentrations of NaCl where applicable. For the NS1 anisotropy experiments, the concentration of RNA or fluorescein labelled NS1-RBD protein was fixed at 10 nM, and the concentration of either NS1 or PABP1 was titrated from 0 to 5 μM. Note for the PABP1 binding to fluorescein labelled poly(A)_18_ experiments, poly(A)_18_ concentration was fixed at 1 nM. Samples were incubated for 1 h at room temperature and the anisotropy studies were performed using a Tecan Spark plate reader in a 96-well plate. The sample (final volume of 100 μL) was excited at 470 nm, and the polarized emission at 520 nm was measured with 10 nm band slits for both excitation and emission. The G-factor was determined using a control sample with fluorescein-labeled RNA. The anisotropy values were subtracted from their initial value, plotted, and fit to the following quadratic equation to determine K_D_ as described previously^21,22^:

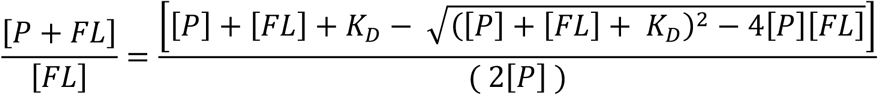

where [P+FL]/[FL] is the anisotropy value, [FL] is the fluorescently labeled species and [P] is the protein concentration. GraphPad Prism (GraphPad Software Inc.) was used to perform the curve fits. All experiments were performed a minimum of three times with different protein batches to ensure reproducibility.

### Purification of Human PABP1

Human PABP1 (GenBank accession code BC015958) in the pANT7_cGST vector was purchased from DNASU. The PABP1 gene was subcloned into pMCSG26, which contains a C-terminal six-His tag.^23,24^ pMCSG26-PABP1 constructs were transformed into *E. coli* Rosetta 2 (DE3) pLysS cells (Millipore). The cells were grown at 37 °C in LB/ampicillin/chloramphenicol to an OD_600_ of 0.6−0.8, cooled to 18 °C, and then induced with 0.25 mM isopropyl β-D-1-thiogalactopyranoside for 12−18 h. Cells were pelleted, flash-frozen, and stored at −80 °C. Cells were resuspended in PABP1 lysis buffer [25 mM Tris (pH 7.5), 250 mM NaCl, 10% glycerol, 8mM DTT, 0.5 mM EDTA, 1 mM PMSF, 0.1% Triton X-100, and 5 mM imidazole] and disrupted by sonication. The cell lysate was centrifuged at 20000g for 45 min at 4 °C. The supernatant was incubated with 4 mL of Ni-NTA beads for 15 min at 4 °C on a rotator. The slurry was poured over a column and washed with 50 mL of PABP1 wash buffer (lysis buffer with 20 mM imidazole and 1mg/mL Heparin sodium salt from porcine intestinal mucosa (Sigma)). Protein was eluted with PABP1 elution buffer [25 mM Tris (pH 7.5), 250 mM NaCl, 10% glycerol, 8 mM DTT, 0.5 mM EDTA, and 250 mM imidazole]. Fractions were collected and concentrated using a 50K MWCO concentrator until the volume was 1 mL. The protein sample was filtered and further purified using a Superdex 16/60 200 pg column at a flow rate of 1 mL/min using PABP1 storage buffer [25 mM Tris (pH 7.5), 250 mM NaCl, 5% glycerol, and 0.25 mM TCEP]. Sample peaks were collected and analyzed by 10% SDS−PAGE. Fractions free of nucleic acids, based on A_280_/A_260_ measurements, were pooled and concentrated using a 50K MWCO concentrator, aliquoted, and flash-frozen. Concentrations of purified proteins were determined by the Bradford assay (Bio-Rad).

### Purification of GST Protein

Plasmid pGEX-3X was used to overexpress and purify the GST protein from *E. coli* BL21 cells as described previously.^13^ The protein was purified using glutathione-Sepharose 4B beads (GE Healthcare) concentrated and flash-frozen in GST buffer [25 mM Tris (pH 7.5), 25 mM NaCl, 10% glycerol, and 0.25 mM TCEP or 0.5 mM DTT] were identified by analyzing aliquots by 12% SDS−PAGE.

### Fluorescence labelling

NS1-RBD FL was labeled with *N*-(5-Fluoresceinyl) maleimide (Sigma-Aldrich) or cyanine 5 (Cy5) maleimide (GE Bioscience), according to the manufacturer’s instructions and as described previously.^25^ Labelled protein was separated from free dye by FPLC on a HiPrep 26/10 Desalting column (Sigma-Aldrich) with NS1-RBD Buffer. Fractions were collected and analyzed by 16% Tricine−PAGE scanned on a FLA9500 Typhoon instrument using the Cy2 or Cy5 excitation laser at a 600 PMT voltage and 50 μm resolution and stained with Coomassie. Fractions free of dye were pooled and concentrated using a 3K MWCO concentrator, aliquoted, and flash-frozen. Concentrations and labelling efficiency of proteins were determined by the Bradford assay (Bio-Rad) and nanodrop.

## Results

### Design of an NS1 RBD mutant that does not form a homodimer

Studies of NS1 have elucidated its ability to dimerize and oligomerize as part of its many functions.^7,26^ Examples include RBD – RBD dimerization to bind directly to RNA as well as ED – ED interactions for the binding to CPSF30 (Figure 1A).^27–29^ We purified the full-length NS1 and the RNA binding domain of NS1 (WT NS1-RBD) because previous studies using pull-down assays showed that the RBD is sufficient for binding to PABP1.^9^ Additionally, the NS1 RBD only forms homodimers, which makes it easier study. To characterize the minimum oligomerization state NS1 must adopt to interact with PABP1, we designed a mutant NS1-RBD that cannot form a homodimer. Previous studies have shown that alanine substitution at Arg 35 and Arg 46 of NS1 disrupts the RBD – RBD interaction.^28^ These mutations were incorporated into an NS1-RBD construct (MUT NS1-RBD), and the protein was purified along with the WT NS1-RBD (Figure 1B). To determine the oligomeric states of the wild type and mutant NS1 RBDs, we analyzed the intrinsic tryptophan-fluorescence polarization of both proteins when they are excited with light at 295 nm wavelength.^30–32^ Based on reports regarding fluorescence polarization of proteins, we expect to see a dimer of NS1 RBD (∼ 16.8 kDa) to polarize light more than the monomer (∼ 8.4 kDa). As predicted, WT NS1-RBD polarizes light more than MUT NS1-RBD, with the difference being in agreement with reports on UV-fluorescence polarization of proteins (Figure 1C).^33^

**Figure 1.**
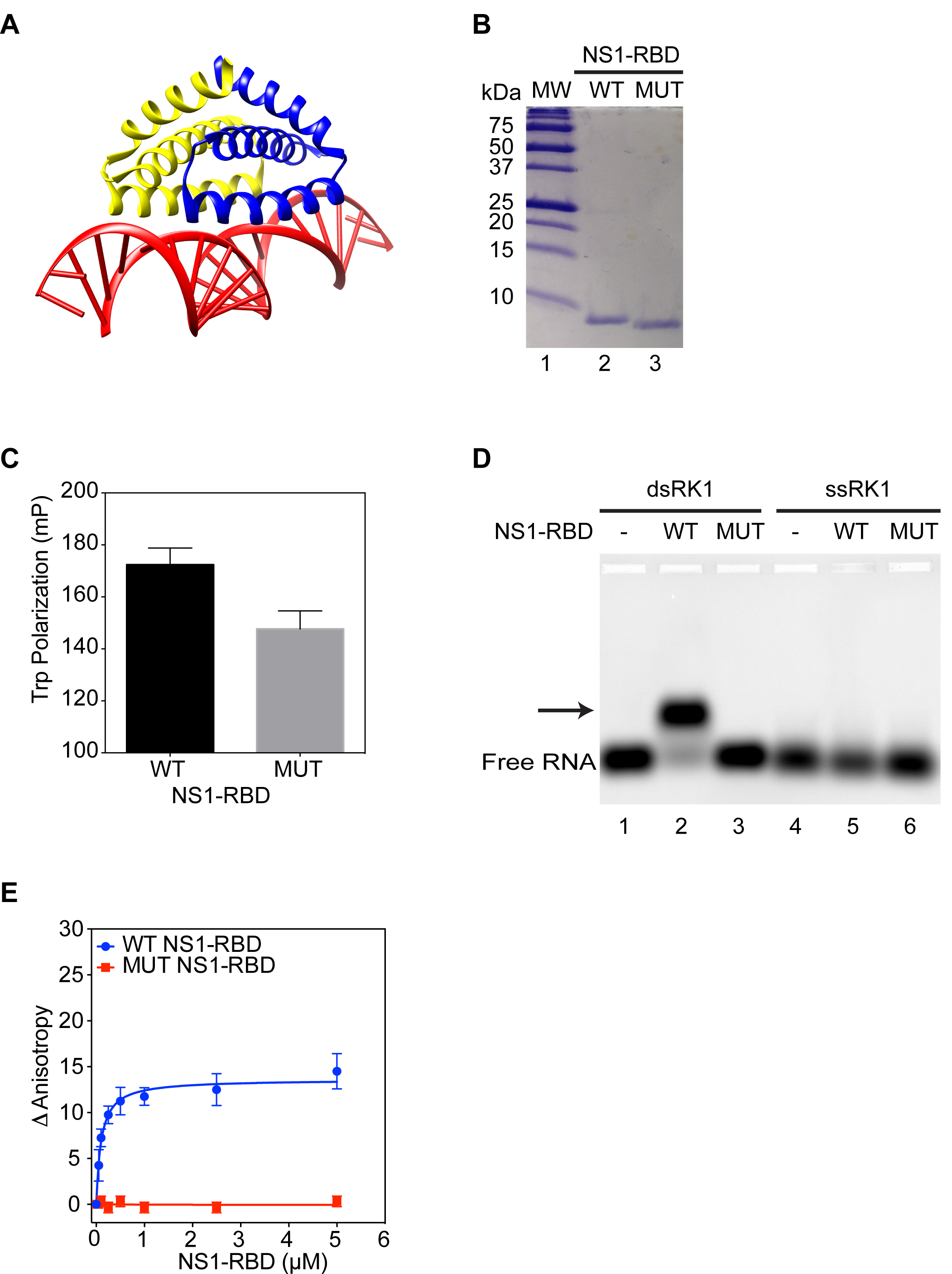
NS1 binds to PABP1 as a dimer. (A) Structure of NS1 RBD dimer (blue and yellow) bound to dsRNA (red) (PDB:2ZKO).^29^ (B) Tricine-PAGE gel of purified WT NS1 RBD and MUT NS1 RBD proteins. (C) Tryptophan polarization of WT NS1-RBD and MUT NS1-RBD. (D) EMSA assay comparing binding of WT NS1 RBD and MUT NS1-RBD to single-stranded RK1 (ssRK1) and double stranded RK1 (dsRK1) RNA. Minus sign indicates no protein was added to the lane. Arrow points to the shifted protein-RNA complex. (E) Anisotropy assay of WT NS1-RBD and MUT NS1-RBD binding to dsRK1 RNA. The final concentration of the RNAs was 10 nM, and the final concentration of NS1 was increased from 0 to 5 μM. The change in anisotropy is shown on the y-axis. The error bars represent the standard deviation from three independent experiments.

Another method for testing the oligomeric states of the NS1-RBD constructs is the ability to bind double-stranded RNA (dsRNA). Based on previous reports that NS1-RBD must be a dimer to bind to dsRNA, we tested the binding of WT NS1-RBD and MUT NS1-RBD to a double-stranded RNA (dsRK1) using an electrophoretic mobility shift assay (EMSA).^27^ We used one of the single-stranded RK1 (ssRK1) sequence as a negative control for the assay. The WT NS1-RBD bound to dsRK1 and slowed its migration on the gel. In contrast, the MUT NS1-RBD did not affect the migration of dsRK1, indicating that the MUT NS1-RBD cannot dimerize to bind to dsRNA (Figure 1D). Additionally, both RBDs do not bind to ssRK1, which is consistent with the specificity of NS1 for dsRNA and not for ssRNA (Figure 1D).

To validate these results, we used a fluorescence anisotropy-based quantitative assay to analyze the binding of NS1-RBDs to dsRK1. Binding experiments were performed by incubating increasing concentrations of NS1-RBD with a fixed concentration of fluorescein-labeled dsRK1 (10 nM), and the change in anisotropy was measured. WT NS1-RBD bound to dsRK1 with a K_D_ of 76 nM ± 9 nM, which is stronger than what has been previously reported for the full-length NS1 (Figure 1E).^13,21^ MUT NS1-RBD incubated with dsRK1 showed no change in anisotropy, indicating that it cannot form a dimer to bind to the dsRNA. Thus, the lower intrinsic tryptophan-fluorescence polarization value of the MUT NS1-RBD and its inability to bind to dsRNA, taken together, supports the conclusion that the MUT NS1-RBD behaves as a monomer in solution.

### Interaction of NS1 with the PABP1•poly(A) RNA complex

We used EMSA to monitor the binding of PABP1 to poly(A) RNA. Briefly, PABP1 was incubated with a 3’ fluorescein-labeled poly adenosine sequence consisting of 18 nucleotides (poly(A)_18_).^13^ In the presence of increasing concentrations of PABP1, we first observe a shift of poly(A)_18_ to an intermediate position on the gel and at a much higher concentration a shift to a higher position compared to the free poly(A)_18_ (Figure 2A). We interpret the complex that migrates to the intermediate position to be a monomer of PABP1 bound to a poly(A)_18_ and the complex that migrates to the higher position to be a PABP1 dimer bound to a poly(A)_18_ RNA molecule. The poly(A)_18_ is short such that only one PABP1 molecule can directly bind to it, and the second PABP1 molecule binds by weaker protein-protein interaction.^14,34,35^ This interpretation is consistent with the fact that PABP1 monomer binds with a high affinity to poly(A) RNA, whereas PABP1 dimers are formed only at much higher concentrations.^36,37^

**Figure 2.**
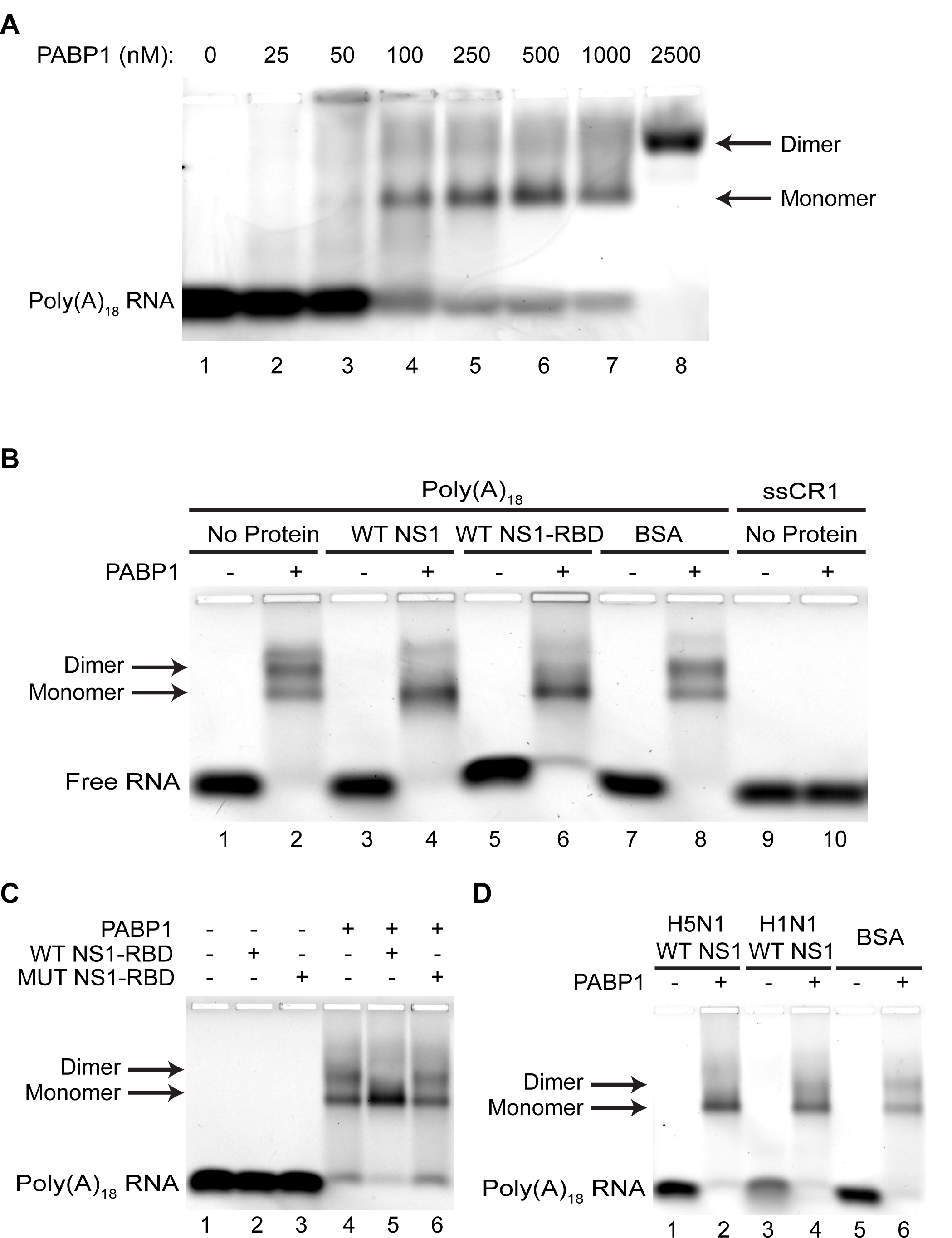
Binding of PABP1 to Poly(A)_18_ RNA in the presence or absence of NS1. (A) PABP1 titration with Poly(A)_18_ RNA. (B) Binding of PABP1 to Poly(A)_18_ RNA in the presence or absence of excess full length H3N2 Udorn NS1 (WT NS1) or the RNA binding domain of H3N2 Udorn NS1 (WT NS1-RBD). BSA serves as a control for the presence of NS1 protein. ssCR1 serves as a control for non-specific binding by PABP1 and NS1. (C) Binding of WT NS1-RBD or MUT NS1-RBD to the PABP1•Poly(A)_18_ complex. (D) Binding of full length H5N1 Nigeria WT NS1 and H1N1 WSN WT NS1 to the PABP1•Poly(A)_18_ complex. Arrows indicate the monomer and dimer of PABP1 on the RNA. Minus and plus signs indicate the absence and presence of NS1, respectively. Arrow points to the shifted protein-RNA complex.

After establishing the EMSA to monitor the binding of PABP1 to poly(A)_18_, we analyzed the effect of NS1 on the PABP1•poly(A)_18_ complex. In the absence of NS1, we observed the two shifted bands corresponding to the PABP1 monomer bound to poly(A)_18_ and the PABP1 dimer bound to poly(A)_18_. Interestingly, in the presence of the full-length NS1, we predominantly observed the band corresponding to the PABP1 monomer bound to poly(A)_18_ (Figure 2B). Similarly, in the presence of the WT NS1-RBD, PABP1 bound as a monomer to poly(A)_18_. These results also agree with previous reports that the RBD is sufficient for binding to PABP1.^9^ As a control, the addition of BSA did not affect the formation of the PABP1 dimer on poly(A)_18_. Additionally, PABP1 does not bind to a control RNA (CR1). These results suggest that NS1, which is present in excess concentration over PABP1, is binding to PABP1 and inhibiting the formation of the PABP1 dimer on poly(A)_18_.

Finally, we analyzed whether the MUT NS1-RBD that is incapable of forming a homodimer affects the binding of PABP1 to poly(A)_18_. Interestingly, in the presence of MUT NS1-RBD, both the PABP1 monomer and the dimer were formed on poly(A)_18_ indicating that the mutant NS1 cannot bind to the PABP1•poly(A) complex, possibly because it cannot homodimerize to form the binding interface needed for interacting with PABP1 (Figure 2C). Thus, NS1 binds as a homodimer to PABP1.

### NS1•PABP1 interaction is conserved across different IAV strains

To date, NS1•PABP1 interaction has been demonstrated only with H3N2 IAV strains, namely A/Victoria/3/75 and A/Udorn/307/1973.^9,13^ Our studies show that in the presence of full-length recombinant A/Udorn/307/1973 NS1, the PABP1 dimer species is disrupted, leaving only the PABP1 monomer bound to poly(A)_18_. To determine whether this interaction occurs with other IAV strains, we purified NS1 proteins from two other strains, namely A/WSN/1933 (H1N1) and A/chicken/Nigeria/2007 (H5N1) (Figure 2D). The NS1 proteins corresponding to the different IAV strains were incubated with the PABP1•poly(A)_18_ complex, and the change in RNA migration was monitored by EMSA. The results show that the different variants of NS1 also disrupt the PABP1 homodimer suggesting that the NS1•PABP1 interaction is conserved across IAV strains. The NS1•PABP1 interaction is conserved across the different IAV strains, most likely due to the RBD’s high sequence homology (Figure S1A).

### Binding of NS1 RBD to PABP1 monitored using a quantitative polarization assay

Although the EMSA results unequivocally show that NS1-RBD binds to PABP1, we wanted a simple, quantitative assay to measure the binding affinity of NS1-RBD for PABP1. Therefore, we decided to develop a fluorescence polarization assay to monitor the binding of NS1-RBD to PABP1. Due to the small size of WT NS1-RBD (∼ 8.4 kDa), we anticipated that a fluorescence polarization assay would be sensitive to the increase in size when NS1-RBD binds to PABP1. WT NS1-RBD has an endogenous cysteine; however, our studies showed that it could not be labeled because it is buried in the protein structure’s interior and is inaccessible to the reactive dye (Figure S1B). Therefore, we added a C-terminal cysteine to WT NS1-RBD (NS1-RBD FL) to label the protein with fluorescein-5-maleimide. Control experiments showed that NS1-RBD with the C-terminal cysteine binds to dsRK1 with the same affinity as WT NS1-RBD, indicating that it is functional (Figure S1C). Additionally, EMSA showed that the NS1-RBD FL decreased the band corresponding to the PABP1 dimer bound to poly(A)_18_, indicating that it is behaving similar to the WT NS1-RBD (Figure S1D).

We next performed the polarization assay to monitor the binding of NS1-RBD FL to PABP1. As predicted, NS1-RBD FL exhibits an increase in anisotropy as the PABP1 concentration increases, showing that the two proteins form a complex (Figure 3A). As a control, we titrated GST protein and did not observe the large increase in anisotropy observed with PABP1. The change in anisotropy with increasing concentrations of PABP1 was plotted, and the data were analyzed by nonlinear regression to obtain a K_D_ of 349 nM ± 69 nM (Figure 3A). The affinity of NS1-RBD for PABP1 is much weaker than the previously reported K_D_ = ∼20 nM between the full-length NS1 and PABP1, which suggests that the ED also contributes to the binding without being necessary.^13^ Furthermore, this weaker binding is consistent with the Nieto group’s qualitative results, where the signal of NS1-RBD used in their pull-down was visibly weaker compared to the full-length NS1.^9^ The new polarization assay can be used to determine the binding affinity of NS1-RBD for PABP1 and could also be used for identifying small molecules that inhibit this interaction.

**Figure 3.**
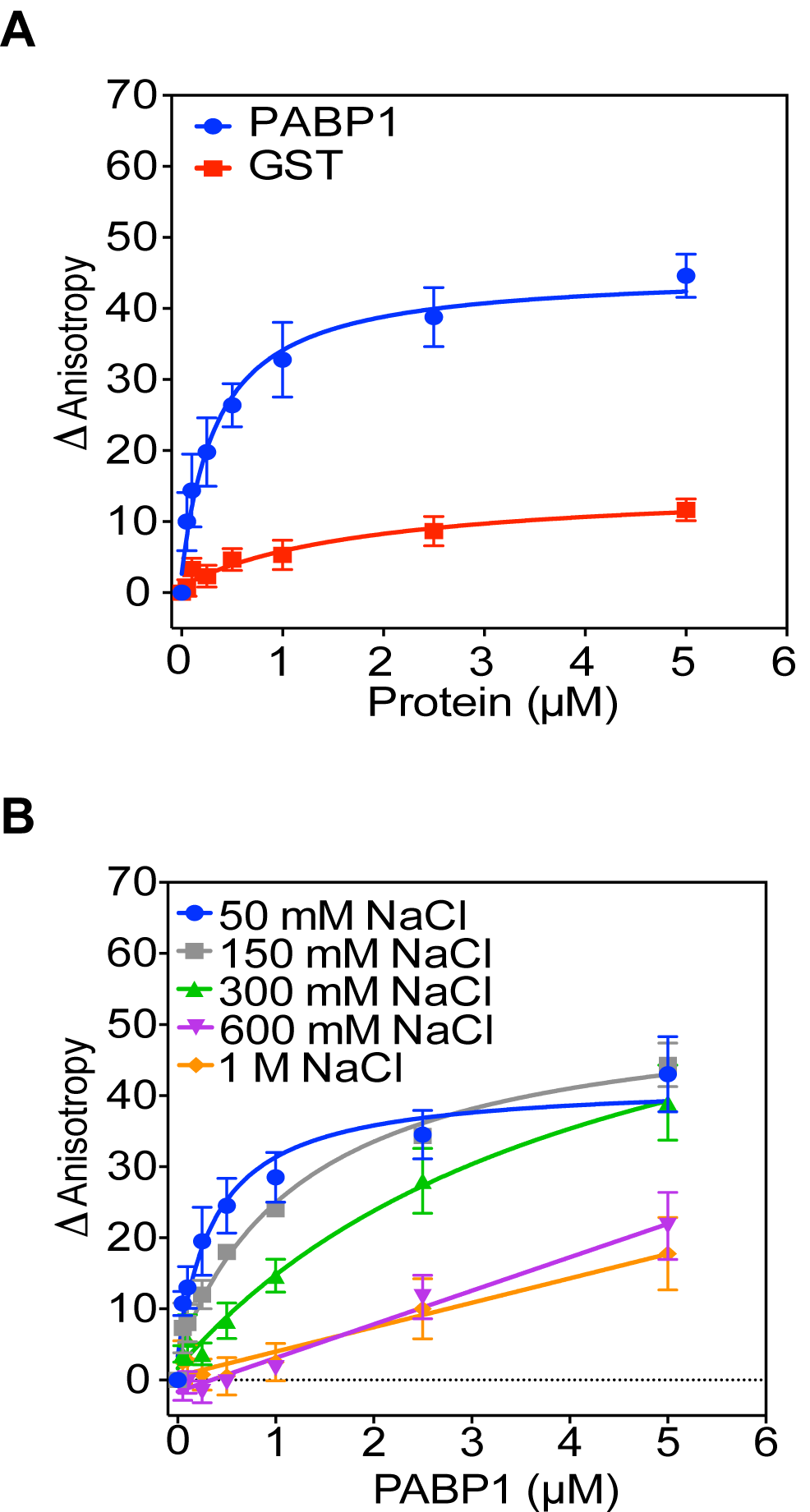
Binding of PABP1 to NS1 RBD monitored with a polarization assay. (A) Plot showing the change in anisotropy when NS1-RBD FL binds to PABP1 or GST as a control. (B) Binding of NS1-RBD FL to PABP1 in the presence of increasing concentrations of NaCl. The final concentration of NaCl ranged from 50 mM to 1M. The final concentration of the NS1-RBD FL was 10 nM, and the final concentration of PABP1 was increased from 0 to 5 μM. The change in anisotropy is shown on the y-axis. The error bars represent the standard deviation from three independent experiments.

NS1 binds to the PABP1 homodimerization domain, which contains interspersed proline residues, polar residues, and positively charged residues that are hallmarks of one type of intrinsically disordered proteins.^38^ We hypothesized that if the interactions were electrostatic, binding of NS1 and PABP1 should weaken as the salt concentration increases. We used the polarization assay to determine the affinity of NS1-RBD FL for PABP1 with increasing concentrations of NaCl to see how sensitive the binding was to competing ions. We found that the affinity of NS1•PABP1 weakened with increasing salt concentrations. For example, the affinity between NS1-RBD FL and PABP1 changed from ∼350 nM at 50 mM NaCl to greater than 1 µM at 150 mM NaCl (Figure 3B). This suggests that NS1 is using the positively charged residues in the homodimerization domain to bind to PABP1.

### NS1 does not bind to the PABP1•poly(A) RNA complex

The EMSA assays’ results made us curious as to where NS1 was migrating on the gel when bound to PABP1. Our studies showed that the addition of NS1 to the PABP1•poly(A)_18_ complex resulted in the disappearance of the band corresponding to the PABP1 dimer bound to poly(A)_18_. Whereas, the PABP1 monomer bound to poly(A)_18_ remained intact. However, we did not detect the potential formation of an NS1•PABP1•poly(A)_18_ complex by EMSA. First, this could be because NS1 binds to the PABP1•poly(A)_18_ complex, and the corresponding change in the isoelectric point of the complex offsets the change in mass giving the appearance of a PABP1 monomer with poly(A)_18_ on the gel. The second possibility is that NS1 cannot bind to the PABP1•poly(A)_18_ complex and is binding only to PABP1 that is free of RNA. However, the formation of the NS1•PABP1 complex cannot be detected because there is no fluorescent dye directly attached to either of the proteins. To resolve these two possibilities, we labeled the NS1-RBD FL construct with the Cy5 dye so that its emission wavelength (670 nm) is distinct from the emission wavelength of the fluorescein dye (521 nm) attached to poly(A)_18_ (Figure S1E). This will allow us to monitor the migration pattern of both NS1-RBD and the PABP1•poly(A) complex simultaneously in the EMSA gels.

We analyzed the binding of the Cy5-labeled NS1-RBD (NS1-RBD-Cy5) to PABP1 and the PABP1•poly(A)_18_ complex by EMSA. Our studies show that the NS1-RBD-Cy5 does not co-localize with the PABP1•poly(A)18 complex but migrates as a smear above and below the PABP1•poly(A)_18_ complex (Figure 4A and B, compare lanes 3, 4, and 6). To verify this observation, we analyzed the binding of NS1-RBD-Cy5 to PABP1 with longer poly(A)_60_ RNA. PABP1 binds to poly(A)_60_ as a dimer that is unaffected in the presence of excess WT NS1 (Figure S2A and S2B).^35,39^ Again, the NS1-RBD-Cy5 does not co-localize with the PABP1•poly(A)_60_ complex but binds to the free PABP1 present in the reaction (Figure 4A and 4B, lanes 9 and 10). Furthermore, we noticed that the NS1-RBD-Cy5 enters the gel only when PABP1 is present (Figure 4A, lanes 5 and 6). To follow up on this observation, we performed a PABP1 titration experiment with a fixed concentration of NS1-RBD-Cy5, which showed that NS1-RBD-Cy5 migrates into the gel only in the presence of PABP1 (Figure 4C).

**Figure 4.**
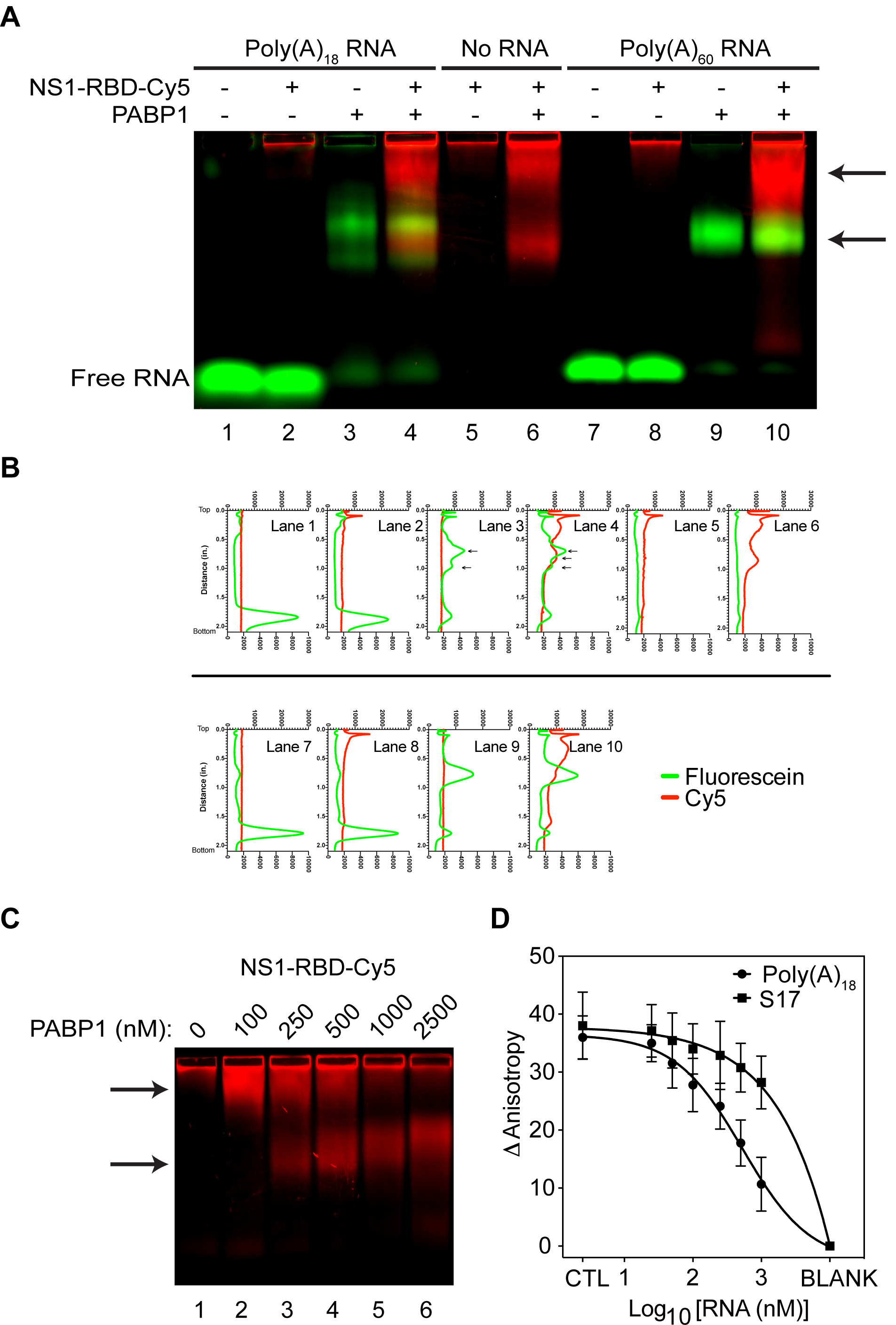
NS1 does not bind to PABP1•Poly(A)_18_ complex. (A) EMSA assay of NS1-RBD labelled with Cy5 (in red) incubated with PABP1 and either Poly(A)_18_ or Poly(A)_60_ RNA labelled with fluorescein (in green). Minus and plus signs indicate the absence and presence of NS1, respectively. Arrows point to the shifted complexes. (B) ImageJ plot of the fluorescence intensity profile for each lane of the EMSA gel in (A). Top axis of each plot represents the intensity (in arbitrary units) of fluorescein (in green) ranging from 0 to 30,000. Bottom axis of each plot represents the intensity (in arbitrary units) of Cy5 (in red) ranging from 0 to 10,000. Left axis of each plot represents relative distance from the well (in inches) ranging from 0 to 2.1. Axes in all plots are scaled the same. (C) EMSA of NS1-RBD binding to varying PABP1 concentrations. NS1-RBD FL is labelled with Cy5. Arrows point to the shifted complexes. (D) Polarization assay with PABP1 pre-bound to NS1-RBD FL (labelled with fluorescein) in the presence of increasing amount of Poly(A)_18_ RNA or S17 RNA. CTL refers to the PABP1•NS1-RBD FL complex in the absence of RNA. BLANK refers to NS1-RBD FL without PABP1 or RNA. The final concentration of the NS1-RBD FL was 10 nM, the final concentration of PABP1 was 500 nM and the final concentration of Poly(A)_18_ RNA and S17 RNA were increased from 0 to 1 μM.

To confirm our results that NS1-RBD binds to the RNA-free PABP1 but not to the PABP1•poly(A) complex, we performed the polarization assay with fixed concentrations of NS1-RBD FL and PABP1 and titrating the concentration of poly(A)_18_. As the concentration of poly(A)_18_ is increased in the reaction, the anisotropy value decreased, indicating that more and more NS1-RBD FL is dissociating from PABP1 because PABP1 is preferentially binding to poly(A)_18_ to form the PABP1•poly(A)_18_ complex. More importantly, the result shows that an NS1•PABP1•poly(A)_18_ complex does not form. A similar experiment with increasing concentrations of the control S17 RNA showed a smaller decrease in anisotropy, consistent with the much lower binding affinity PABP1 has for S17 (Figure S2C). Thus, these results show that NS1 does not bind to the PABP1•poly(A) RNA complex.

## Discussion

The NS1 protein is a multifunctional protein that binds to dsRNA and to several host proteins to enable IAV to replicate efficiently in host cells.^7,40,41^ NS1 can form dimers and oligomers by itself, and a long tube-like multimer that wraps around dsRNA.^42^ Previous studies also showed that NS1 binds as a dimer to TRIM25, CPSF30, DHX30 and dsRNA.^28,43–45^ Thus, depending on the interacting partner, NS1 displays various oligomeric states that are critical for its function.^41^ Here we show that NS1 monomer cannot bind to PABP1, but NS1 that can form dimers bind to PABP1. This is not unexpected because NS1 is expressed to high levels in the infected cells and may exist predominantly as dimers or other higher-order structures.^46,47^

PABP1 is one of the most highly expressed proteins in the cell and plays an essential role during translation initiation.^11,48^ PABP1 binds to the 3’ poly(A) tail of mRNA and protects the mRNA from degradation by exonucleases.^49,50^ Additionally, PABP1 interacts with the eIF4F complex, which is bound to the 5’ 7-methyl guanosine cap structure, to bring the 5’- and 3’-ends close together to form the mRNA closed-loop structure.^51^ The mRNA closed-loop structure is thought to stimulate translation by recycling the terminating ribosome back to the 5’-end of the mRNA to initiate translation.^52^ PABP1 binds to poly(A) with a high affinity (K_D_ ∼ 5 nM) and can bind to a poly(A) sequence as short as 12 nucleotides with no change in binding affinity.^34,35^ Although PABP1 can bind to short poly(A) tails with high affinity using its RRM domains, PABP1 covers about 30 nucleotides because of steric occlusion by the rest of the protein.^14,34,35,39^ Recent studies indicate that most mRNAs have a poly(A) tail that is 30 nt in length,^53^ suggesting that only one PABP1 is directly bound to the mRNA, and possibly a second PABP1 may bind via protein-protein interaction. Our results show that NS1 cannot bind to PABP1 bound to the mRNA poly(A) tail. This suggests that the poly(A) RNA bound to the RRM domains sterically blocks the binding of NS1 to the homodimerization domain of PABP1. Alternatively, NS1 can only bind to the RNA-free PABP1 because it has a different conformation than the PABP1 bound to the poly(A) RNA. Importantly, the function of the NS1•PABP1 complex appears to be distinct from the classical role of PABP1 in translation initiation, when it is bound to the 3’-poly(A) tail of mRNA.

Previously, we showed that NS1 could not bind simultaneously to both dsRNA and PABP1. The RBD of NS1 is responsible for binding to dsRNA and PABP1, and it can accommodate only one of these partners. Here, we show that NS1 binds to PABP1 free of poly(A). Thus, both proteins can interact with each other only when they are not bound to RNA. We expect PABP1 will bind to all the available mRNA poly(A) tails because the affinity of PABP1 for poly(A) is significantly higher than for NS1. However, PABP1 is present in excess over the total cellular mRNA concentration, and it is estimated that only 30% of the PABP1 molecules are bound to the poly (A) tail.^11^ Thus, NS1 can interact with the large pool of PABP1 molecules that are not bound to the poly(A) RNA.

The functional significance of the interaction of NS1 with PABP1 for the life cycle of IAV is unknown. Because NS1 also interacts with eIF4G, it has been suggested that NS1 may stimulate the translation of viral mRNAs by promoting the binding of the 3’ poly(A)•PABP1 complex to the eIF4F complex present at the 5’-end of the mRNA.^9,15,54–56^ However, our results are not consistent with this model because we show that NS1 cannot bind to PABP1 that is bound to poly(A) RNA. Interestingly, PABP1 interacts with the ribosome directly, and this interaction was shown to enhance translation in a dose-dependent manner.^16^ It has been proposed that PABP1 may stimulate translation initiation by promoting the recruitment of the 40S and 60S subunits to the eIF4F initiation complex that is assembled at the 5’-end of the mRNA.^16–18,57,58^ Additionally, NS1 is a general enhancer of translation, and a recent report showed that NS1 stimulates the binding of the ribosome to the mRNA.^59–61^ Taken together, we propose that NS1’s ability to bind to both eIF4G and the RNA-free PABP1 may be a mechanism to recruit PABP1 to the 5’-end of the mRNA. The NS1•eIF4F•PABP1 complex at the 5’-end of the mRNA may then stimulate translation by enhancing the recruitment of the ribosomal subunits to the initiation complex.

## ACCESSION CODES

UniProtKB: P11940 (PABP1), P03495 (H3N2 NS1), D2SJD8 (H5N1 NS1), Q82506 (H1N1 NS1)

## Author Contributions

C.M.D. and S.J. designed the experiments, C.M.D. performed the experiments, C.M.D. and S.J. wrote the manuscript.

## Notes

The authors declare no competing financial interest.

## Supporting Information

Additional NS1 and PABP1 binding data and quality control experiments.

