## Supplemental information for "Influenza A virus NS1 protein binds as a dimer to the RNA-free PABP1 but not to the PABP1•Poly(A) RNA Complex"

La Jolla, CA 92093-0314, USA

* To whom correspondence should be addressed. Tel: + 858 822 2957;

**Supporting Information Figure 1.** NS1 quality control. (A) Primary sequence alignment of the RNA Binding domain of the four NS1 strains used to demonstrate binding to PABP1. Yellow background highlights residues that are identical across all four strains. (B) Tricine-PAGE of WT NS1-RBD and NS1-RBD FL after fluorescein labeling. Gel on left is after Coomassie staining. Gel on right is scanned with the Typhoon FLA9500 using the Cy2 channel. (C) Polarization assay comparing the RNA binding properties of WT NS1-RBD, MUT NS1-RBD, and NS1-RBD FL with dsRK1. The final concentration of the RNA was 10 nM, and the final concentration of NS1 was increased from 0 to 5 μM. The change in anisotropy is shown on the y-axis. The error bars represent the standard deviation from three independent experiments. The $K_D$ for WT NS1-RBD, and NS1-RBD FL binding to dsRK1 are 96 nM ± 19 nM and 115 nM ± 27 nM, respectively. (D) EMSA showing the binding of NS1-RBD FL to PABP1•poly$(A)_{18}$ complex. The concentrations of poly$(A)_{18}$, PABP1, and NS1-RBD FL are 100 nM, 500 nM and 5000 nM, respectively. GST at 5000 nM was used as a control for NS1-RBD FL. (-) and (+) indicate the absence and presence of the protein, respectively. The arrows indicate the PABP1 monomer and dimer bands. (E) Tricine-PAGE of NS1-RBD FL after Cy5 labeling. Gel on left is after Coomassie staining. Gel on right is scanned with the Typhoon FLA9500 using the Cy5 channel.

**Supporting Information Figure 2.** Binding of PABP1 to RNA. (A) EMSA showing the binding of PABP1 to the fluorescein-labeled poly$(A)_{60}$ RNA. The concentration of the RNA is 100 nM and the concentration of PABP1 was titrated, as indicated. Poly$(A)_{18}$ at 100 nM concentration was used as a control. The arrows indicate the PABP1 monomer and dimer bands. (B) Binding of PABP1 to Poly$(A)_{60}$ RNA in the presence or absence of excess full length H3N2 Udorn NS1 (WT NS1). (C) Polarization assay showing the binding of PABP1 to poly$(A)_{18}$ and S17 RNA. The final concentrations of the fluorescein-labeled poly$(A)_{18}$ and S17 RNA were 1 nM and 10 nM respectively, and the final concentration of NS1 was increased from 0 to 5 μM. The change in anisotropy is shown on the y-axis. The error bars represent the standard deviation from three independent experiments.

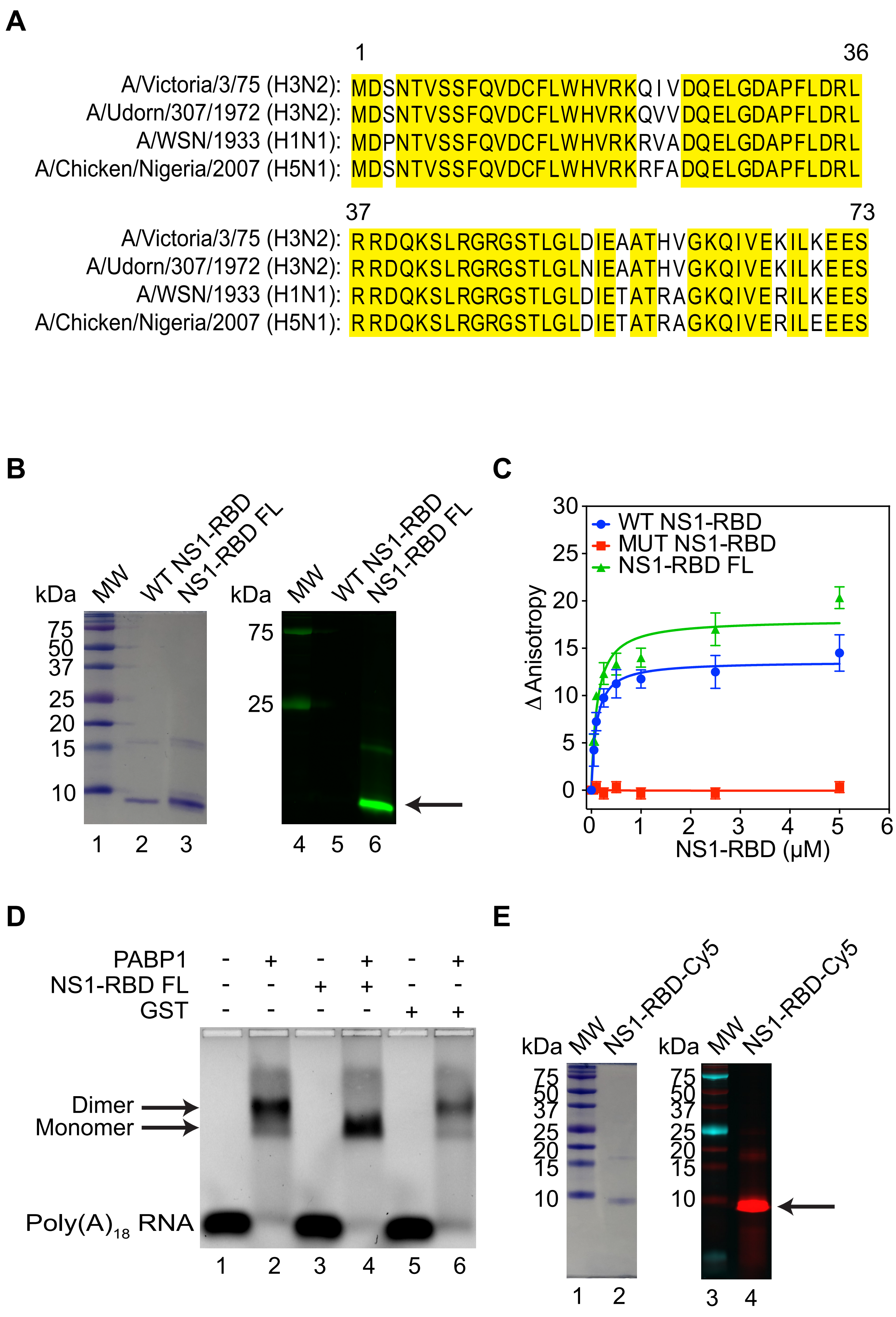


**Supporting Information Figure 1**

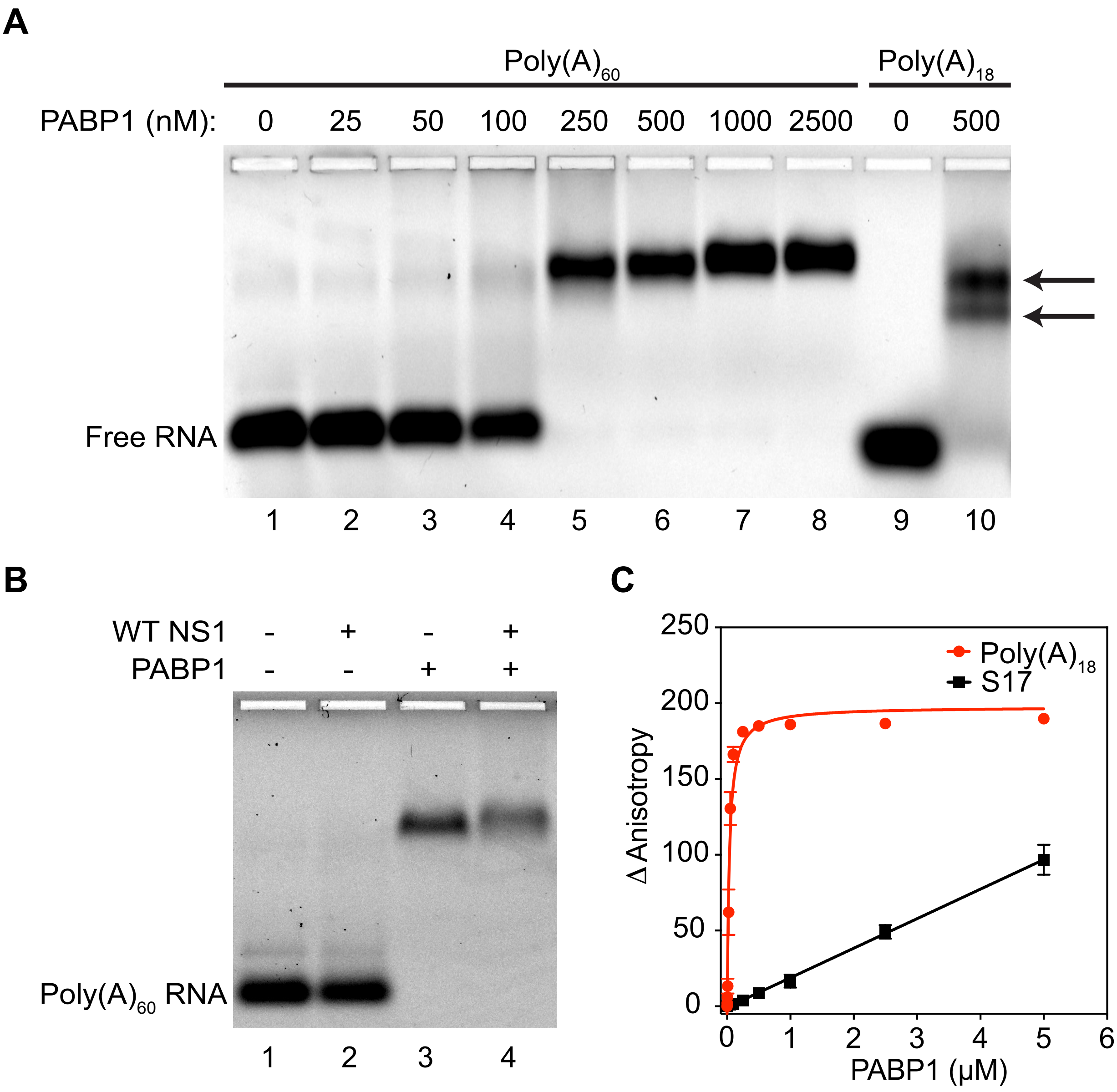


**Supporting Information Figure 2**
